# Glycosylation-enabled Site-specific Growth Factor Engineering for Biomaterial Functionalization

**DOI:** 10.1101/2025.05.11.652418

**Authors:** Yunhui Xing, Qingyang Li, Phil G. Campbell, Xi Ren

## Abstract

In native tissue environments, growth factors (GFs) are often physically associated with the extracellular matrix (ECM) framework. Despite the enormous potential of the chemoselective click chemistry for GF functionalization of biomaterials to recapitulate such GF-ECM association, its application is limited by the lack of a universal strategy for reliable production of clickable GFs. Here we present a novel platform technology that leverages intrinsic post-translational protein glycosylation to enable site-specific metabolic engineering of GFs with azido tags for subsequent ECM hydrogel conjugation. Using Vascular Endothelial Growth Factor as a model, we demonstrated efficient, glycosylation-dependent azido incorporation during its recombinant expression with preserved bioactivity. We further expanded the utility of this strategy to non-glycosylated proteins through engineered N-linked glycosylation via the incorporation of a signal peptide that directs newly synthesized proteins to the secretory pathway where glycosylation takes place along with a sequon for glycan attachment. The resulting GF with site-specific azido incorporation can be effectively immobilized within dibenzocyclooctyne-bearing ECM hydrogel via the copper-free click chemistry, exhibiting sustained GF retention and delivering augmented angiogenic responses. Our approach thereby offers an unprecedented opportunity to streamline recombinant protein engineering for biomaterial functionalization in tissue engineering and regenerative medicine applications.

## 1. Introduction

Growth factors (GFs) are essential mediators of intercellular communication, playing critical and diverse roles in regulating cell proliferation, migration, and differentiation. They induce tissue morphogenesis, support tissue homeostasis, and drive repair and regeneration [1-3]. Upon secretion into the extracellular space, the range and magnitude of GF function largely depend on their interactions with other extracellular matrix (ECM) components, such as fibrous proteins (e.g., collagens and laminins) and glycosaminoglycans (GAGs), which modulate GF diffusivity and stability within tissue microenvironments [4-8]. Frequently, ECM binding provides GFs with resistance to protease digestion, enhances their receptor activation ability, and facilitates localized influence by maintaining above-threshold GF concentrations within a small range from the source [9-13]. Additionally, the ECM can function as a reservoir for triggered GF release in response to matrix remodeling events, thereby providing a spatiotemporally dynamic and responsive signaling environment [7, 14, 15].

Recent advances in biomaterial engineering have introduced a wide range of modified ECM matrices for GF delivery, aimed at enhancing their stability and efficacy. For example, researchers have utilized sulphated chondroitin sulfate glycosaminoglycan hydrogel, heparin-functionalized PEG gel, or the combined use of GAGs such as dermatan sulfate, heparin, and hyaluronic acid hydrogel to deliver various bioactive substances, including fibroblast growth factor (FGF), bone morphogenetic protein-2 (BMP-2), and interleukin [1-7, 9]. While achieving long-term retention, they successfully maintained the locally enriched the release of substances and ensured the corresponding functions.

Conventional strategies for GF-to-biomaterial immobilization typically utilize amine- or thio-reactive chemistry that targets the lysine and cysteine residues on the GFs. Given the high prevalence of these amino acid residues and their frequent involvement in GF function, such conjugation reactions must be individually assessed and optimized based on the amino acid sequences of each GF of interest, which limits their broad applicability. Therefore, a platform technology that enables site-specific GF modification is highly desired, as it will streamline the GF-to-biomaterial conjugation process while preserving the bioactivity of GFs. Chemoselective chemistry, such as azide-to-alkyne cycloaddition, also known as click chemistry, presents a promising alternative by facilitating highly specific binding between biologically inert ligands [16-21]. Nevertheless, a universal method to render GFs clickable has not yet emerged.

All GFs are initially synthesized and secreted by cells, with glycosylation being one of the most common post-translational modifications to secreted proteins, including GFs. This process is directed by signal peptides, which are short sequences of amino acids that guide proteins into the endoplasmic reticulum (ER) for glycosylation, where glycan chains attach to specific amino acid motifs known as sequons on the newly synthesized protein [22, 23]. Previous research from us and others has shown that bioorthogonal glycoprotein engineering using azido-monosaccharide probes facilitates the incorporation of click-reactive azido tags into the glycan chains of newly synthesized ECM proteins, allowing for subsequent chemoselective protein modification and conjugation[20, 24]. Although most proteins can be glycosylated either naturally or artificially, there have been limited attempts to leverage glycosylation for GF engineering.

In this study, we developed a platform technology that metabolically labels engineered GFs with azido tags and immobilizes them onto click-reactive ECM-derived hydrogels for controlled GF delivery. We utilized vascular endothelial growth factor 165 (VEGF165), a glycosylated growth factor, to demonstrate our labeling approach, given its widespread application in promoting vascularization in tissue engineering [1, 2, 6, 10, 25-27]. To extend the platform to non-glycosylated proteins, we also engineered a glycosylated enhanced green fluorescent protein (eGFP) by incorporating a signal peptide and sequons, enabling bioorthogonal azido tag incorporation into the newly synthesized glyco-eGFP. In parallel, we functionalized collagen and fibrin hydrogels with azido-reactive dibenzocyclooctyne (DBCO) and validated efficient and selective covalent conjugation between azido-GFs and DBCO-ECM hydrogels via the copper-free click chemistry. The resulting GF-functionalized hydrogels exhibited stable long-term retention of the immobilized GFs and delivered enhanced pro-angiogenic functions in vitro.

## 2. Material and methods

### 1. Plasmid design and cell culture

The coding sequence of human Vascular Endothelial Growth Factor 165 (VEGF165) was fused with a C-terminal His-tag (Table S1) and cloned into the pCDH-CMV-MCS-EF1-Puro lentiviral vector (System Biology, CD510B-1), referred to as the VEGF165 lentiviral vector. The eGFP coding sequence was engineered to include combinations of signal peptides and glycosylation Sequon, and cloned into the pcDNA 3.1(+) Mammalian Expression Vector (Invitrogen), referred to as the engineered eGFP vectors, for transient expression. Signal peptides used were from either human VEGF165 (SP_VEGF_) or the rat Follicle-Stimulating Hormone β-subunit (SP_FSH_), and the glycosylation Sequon was derived from VEGF165 (Sequon_VEGF_) (Table S1). Cloning was performed by GenScript Biotech Co., Ltd.

Chinese hamster ovary (CHO) cells (ATCC, CCL-61) were maintained in DMEM/F12 with 10% Fetal Bovine Serum (FBS). 293T cells (ATCC, CRL3216) were cultured in DMEM with 10% FBS. Human umbilical vein endothelial cells (HUVECs) (Lonza, C2517A) were maintained in Endothelial Cell Growth Medium-2 (EGM-2, Lonza, CC-3162). The stable VEGF165-overexpressing CHO cell line was established with the third-generation lentiviral transfection. Briefly, the VEGF165 lentivirus was packaged and harvested from 293T cells that were co-transfected with pRSV-REV, pMDLg/pRRE, pMD2.G, and the VEGF165 lentiviral vector. The CHO cells were then transduced by the harvested VEGF165 lentivirus-conditioned medium supplemented with 8 µg/mL polybrene (MilliporeSigma, TR1003G) and selected with 10 µg/mL puromycin (Gibco, A1113802).

### 2. Metabolic glycan labeling and purification of VEGF165

For metabolic glycan labeling of VEGF165, the VEGF165-overexpressing CHO cells were cultured with 50 µM of Ac4ManNAz (Click Chemistry Tools, CCT-1084) or 0.1% DMSO (vehicle control) for 24 h prior to switching to the collection medium (DMEM/F12 with 1% GlutaMAX (Gibco, 35050061) and MEM non-essential amino acids (Gibco, 11140050)) supplemented with the same labeling reagents. Following another 72 h of culture, the resulting VEGF165 conditioned medium was collected, centrifuged at 2,000× g for 10 min at 4°C to remove the cell debris, and concentrated to 1/10 of the initial volume with 10,000 MWCO ultrafiltration filters (Millipore Sigma, UFC901008). VEGF165 was then affinity-purified from the concentrated conditioned medium using HIS-Select® Nickel Affinity Gel (Sigma, P6611). The purified VEGF165 was either used immediately or stored at -80 °C. The VEGF165 concentration was determined using the VEGF-A Human ELISA Kit (Invitrogen, BMS277). The VEGF165 purified from culture with and without Ac4ManNAz supplementation was referred to as Az-VEGF and non-Az-VEGF, respectively. To perform the click shift assay, the Az-VEGF/non-Az-VEGF was click-conjugated with 20 µM DBCO-PEG, 30 kDa (Click Chemistry Tools, CCT-A121) at room temperature (RT) overnight and then analyzed by gel electrophoresis and Western blot.

### 3. Bioactivity assays of glycan-engineered VEGF165

For endothelial metabolic activity assay, 3,000 HUVECs were seeded into each well of the 96-well microplate and cultured overnight in EGM-2. Medium was then switched to the serum- and supplement-free Endothelial Cell Basal Medium-2 (EBM-2, Lonza, CC-3156) with or without Az-VEGF/non-Az-VEGF at a series of concentrations of 5, 10, or 25 ng/mL. Positive control culture was continued in fully supplemented EGM-2. These cultures were continued for another 24 h prior to metabolic assay in EBM-2 with 50 µM of resazurin for 2 h at 37°C in dark. The relative fluorescence intensity (Ex 544 nm, Em 590 nm) of the resulting medium was quantified using the SpectraMax microplate reader (Molecular Devices).

Endothelial transwell chemotaxis assay was performed in 24-well microplates with polycarbonate transwell inserts with 8-µm pore size (Millipore-Sigma, ECM508). HUVECs were seeded onto the upper chamber of the transwell insert at 50,000 cells per insert in the basic EBM-2 (with 0.5% FBS). 600 µL of EBM-2 (with 0.5% FBS) with or without Az-VEGF/non-Az-VEGF (5, 10, or 25 ng/mL) was added to the lower chamber. For the positive control culture, 600 µL of fully supplemented EGM-2 was added to the lower chamber. Following 24 h of incubation, cells on the insert’s top surface (non-migrated cells) were removed using a cotton swab, and then the insert together with cells that have migrated through the pores to its bottom surface was fixed with ice-cold methanol for 10 min at 4°C and stained with 0.2% crystal violet for 10 min at RT. Three random bright-field images were taken per insert using a 20X objective of EVOS M5000 microscope. The extent of cell migration per condition was represented as the percentage of migrated cell coverage within each image.

### 4. DBCO modification of fibrinogen

To prepare DBCO-modified fibrinogen (DBCO-fibrinogen), plasminogen-depleted bovine fibrinogen (Enzyme Research Laboratories, BFIB1) was diluted in PBS to 10 mg/mL, and incubated with 200 µM DBCO-N-hydroxysuccinimidyl (DBCO-NHS) ester (Click Chemistry Tools, A133) in 10% DMSO for 3 h at RT. The reaction mixture was then extensively dialyzed against PBS using the 100 kDa MWCO ultrafiltration filter (Millipore Sigma, UFC9100) for removal of unconjugated DBCO and buffer exchange.

### 5. Gel electrophoresis, Western blot and dot blot

Protein samples were separated in 4-15% sodium dodecyl sulfate-polyacrylamide gel electrophoresis (SDS-PAGE) gels. For on-gel protein staining, SYPRO Ruby Protein Gel Stains (Invitrogen, S12000) was used. For Western blot, following transfer onto polyvinylidene fluoride (PVDF) membranes (ThermoFisher Scientific, 88518), blots were incubated overnight at 4 °C with anti-His-tag antibody (1:500; Santa Cruz, sc-53073) or anti-GFP antibody (1:500; Santa Cruz, sc-9996) and subsequently for 1 h at RT with the horseradish peroxidase (HRP)-conjugated secondary antibody (1:10,000; Pierce, SA1100, ThermoFisher Scientific), and then blots were developed with SuperSignal™ West Pico PLUS Chemiluminescent Substrate (ThermoFisher Scientific, 34580).

To verify DBCO modification of fibrinogen, the unmodified fibrinogen and DBCO-fibrinogen were diluted to 10 mg/mL with PBS, click-conjugated with 50 µM Azide-Cy5 (Az-Cy5, Click Chemistry Tools, AZ118) for 1 h at RT, and extensively dialyzed against PBS using the 3 kDa MWCO ultrafiltration filter. To perform dot blot analysis, the dialyzed reaction mixtures were applied onto nitrocellulose membrane, air dried, washed with tris-buffered saline with Tween-20 (TBST), and imaged with the ChemiDoc Imaging System (Bio-Rad). In parallel, total protein loading was assessed by on-blot staining with the Ponceau S solution (Sigma Aldrich, P7170).

### 6. Metabolic glycan labeling and deglycosylation of engineered eGFP

CHO cells were transfected with vectors expressing native and engineered eGFP utilizing the Lipofectamine 3000 Transfection Reagent (Invitrogen, L3000008) in collection medium with 50 µM of Ac4ManNAz or 0.1% DMSO. To assess the subcellular localization of the expressed native and engineered eGFP, 24 h following transfection, cells were imaged with a 20X objective of the EVOS M5000 fluorescent microscope. Conditioned medium was collected from the transfected cells 48-72 hours following transfection, centrifuged at 2,000x g to remove cell debris, and extensively dialyzed against PBS using the 3 kDa MWCO ultrafiltration filter (Millipore Sigma, UFC5003BK) to remove free Ac4ManNAz. For click shift assay, conditioned medium containing native or engineered eGFP was clicked with 20 µM DBCO-PEG, 30 kDa at RT overnight, and then analyzed by Western blot. Deglycosylation of engineered eGFP was performed via treatment of PNGase F (New England Biolabs, P0704S) for 1 h at 37°C prior to click-conjugation with DBCO-PEG, 30 kDa.

### 7. In vitro retention of Az-Cy5 in fibrin hydrogel

Unmodified fibrinogen or DBCO-fibrinogen (15 mg/mL) was mixed with 10 U/mL of bovine thrombin (Millipore Sigma, 6051571KU) together with Az-Cy5, transferred to 96-well microplate (50 µL per well), and incubated for 1 h at 37 °C to allow gelation. 100 µL of PBS was then added to each fibrin gel well and refreshed daily to remove the released fluorophore. Fluorescence intensity of each fibrin gel was measured daily following each PBS exchange using the SpectraMax microplate reader.

### 8. Radiolabeling VEGF165 with ^125^I

VEGF165 was iodinated in a polypropylene 12×75 mm test tube using the Iodogen bead method. 5 µg VEGF165 was reacted with 400 µCi ^125^I-Na in PBS for 30 min at 25 °C in the presence of 2 iodogen beads (Fisher Scientific) with occasional swirling. At 30 min, the reactant liquid was transferred to a Sephadex G25 column and 0.5 mL fractions were eluted using PBS. The VEGF165 fraction was aliquoted and stored at -80 °C.

^125^I-labeled VEGF165 (Az-VEGF or non-Az-VEGF) of approximately 106 counts per min (cpm) radioactivity was mixed with 300 µL of the pre-gel mixture for unmodified fibrin or DBCO-fibrin hydrogels. Release kinetics was assessed in simulated body fluid (SBF: 10 % FBS, 0.02 % sodium azide, 25 mM HEPES in DMEM) as described previously [28]. Briefly, gels were placed in 12 × 75 mm polypropylene tubes each containing 1 mL of SBF. Tubes were incubated at 37 °C for the indicated time durations, SFB was then replaced and the retained radioactivity was measured using the Wizard2 2-Detector Gamma Counter (PerkinElmer, Waltham).

### 9. 3D endothelial metabolic activity assay with hydrogel

A double-layer hydrogel system was established, with the bottom-layer hydrogel containing VEGF-functionalized collagen gel and the top-layer hydrogel embedded with endothelial cells. For the bottom-layer hydrogel, Az- or non-Az-VEGF at varying concentrations of 0, 50, or 200 ng/mL was mixed with 2 mg/mL DBCO-fibrinogen (or control unmodified fibrinogen) and 1 U/mL thrombin in PBS to prepare the fibrin gel. A 50 µL pre-gel mixture with Az- or non-Az-VEGF was seeded into each well of the 96-well plate and incubated at 37 °C for 10 min to allow gelation before adding 200 µL of PBS on top of the gel. PBS was replaced 10 times within 48 h to facilitate the release of non-conjugated VEGF. For the top-layer hydrogel, 10,000 HUVECs were resuspended in 50 µL of pre-gel mixture containing 2 mg/mL unmodified fibrinogen and 1 U/mL thrombin, then rapidly inoculated onto the pre-washed fibrin hydrogel (with and without click-conjugated VEGF as described above). One hundred µL of EBM-2 (with 0.5% FBS) was added on top of the double-layer hydrogel system, and the culture proceeded for 48 h before the resazurin metabolism assay in EBM-2 with 50 µM of resazurin for 4 h at 37°C in the dark. Positive and negative control cultures were prepared using a similar double-layer hydrogel system but with unmodified fibrin hydrogel without VEGF as the bottom-layer hydrogel and with EGM-2 or EBM-2 (with 0.5% FBS) as the culture medium, respectively.

### 10. Endothelial scratch recovery assay with hydrogel

Fibrin groups under testing were prepared and washed as described in Section 2.9, with 50 µL of hydrogel placed on top of each transwell insert featuring an 8-µm pore size (cellQART, 9328002) within the 24-well plate. In parallel, 50,000 HUVECs were seeded per well in the 24-well plate and cultured in EGM-2 for 24 h. A 2-mm-wide scratch was created with a cell scraper (VWR, 76036-006) in each well, after which the hydrogel-containing transwell insert described above was positioned on top of the HUVEC culture with the scratch. EBM-2 (with 0.5% FBS) was added to both the upper and lower chambers, and the culture continued for another 24 h to enable scratch recovery. The scratch was imaged under bright field with a 10X objective of the EVOS M5000 microscope both before and after the recovery phase, and its width was measured to calculate the percentage of scratch recovery.

### 11. Endothelial transwell chemotaxis assay with hydrogel

Fibrin groups under testing were prepared and washed as described in Section 2.9, using 75 µL of hydrogel per well of the 24-well plate. A transwell insert with an 8-µm pore size (cellQART, 9328002) was placed on top of each well, and 50,000 HUVECs in 150 µL of EBM-2 (with 0.5% FBS) were seeded onto each insert in EBM-2 (with 0.5% FBS). Additionally, 600 µL of EBM-2 (with 0.5% FBS) was added to the lower chamber above the pre-washed fibrin hydrogels. For the positive control culture, the fibrin hydrogel without VEGF and fully supplemented EGM-2 was utilized. The remainder of the transwell chemotaxis assay was performed and analyzed as described in Section 2.3.

### 12. Statistics

Quantitative data were displayed as the mean ± standard error of the mean (SEM). Statistical comparisons were performed using one-way analysis of variance (ANOVA) with post-hoc Tukey’s test and unpaired two-tailed Student’s t-test. Statistical analyses were performed using GraphPad. * *p*<0.05, ** *p*< 0.01, *** *p*<0.001.

### 13. Graphics

Schematics were created using Microsoft PowerPoint and BioRender. Plots were prepared using GraphPad. Chemical structures were prepared using MarvinSketch.

## 3. Results

### 3.1 Metabolic glycan engineering of human recombinant VEGF165

We developed a strategy to integrate a metabolic glycan labeling process into the recombinant expression of human VEGF165 glycoproteins, enabling site-specific incorporation of the azido (Az) tag onto its glycan chain, referred to as Az-VEGF. The resulting Az-VEGF allows for chemoselective immobilization into extracellular matrix (ECM) hydrogel bearing the complementary DBCO modification through the copper-free azide-to-DBCO click reaction, to control VEGF retention and release (Figure 1A).

**Figure 1.**
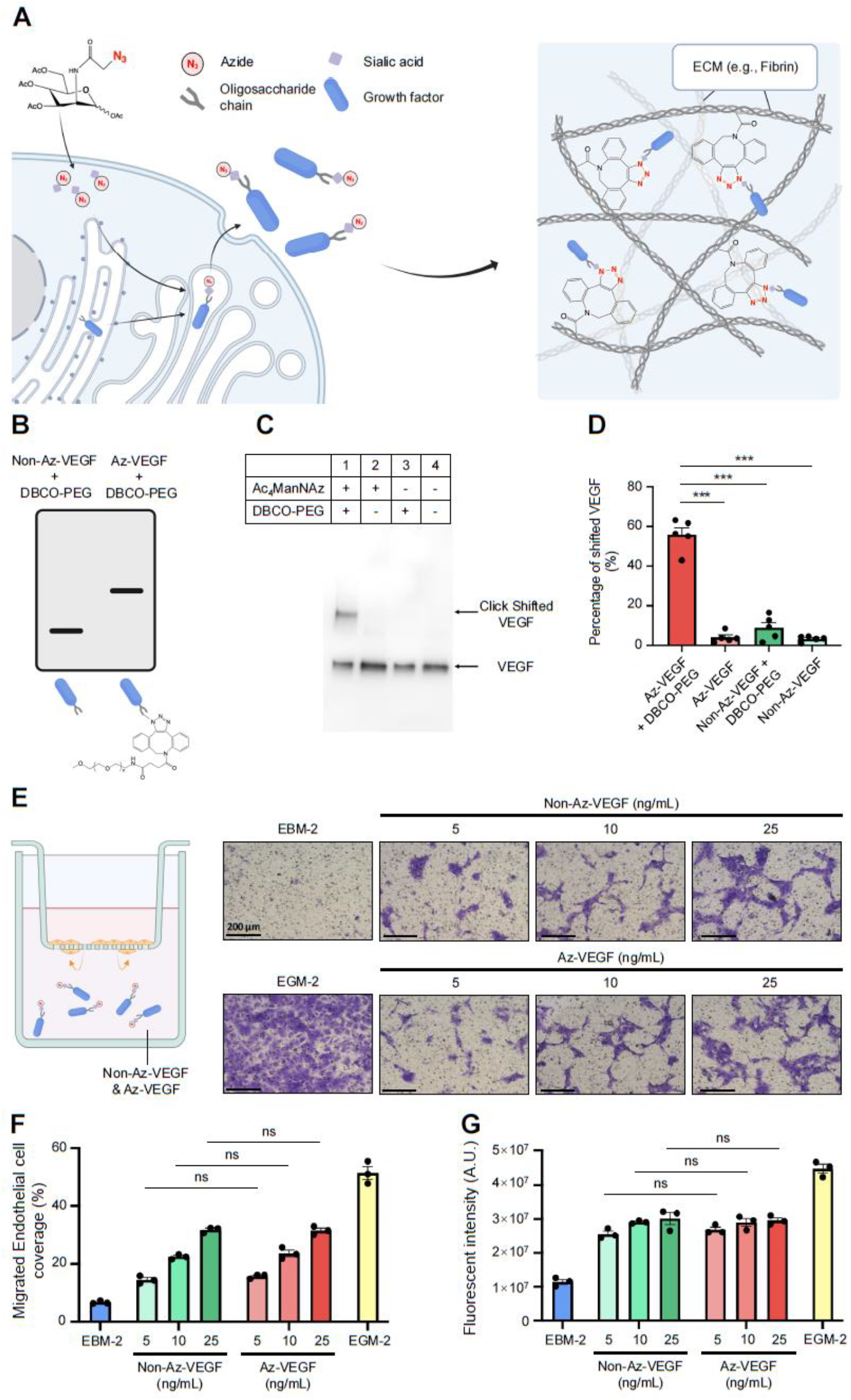
Metabolic glycan engineering of human recombinant VEGF165. (A) Schematic showing the overall strategy for metabolic glycan engineering GF to incorporate azido labeling. Created with BioRender.com. (B-D) VEGF165 click shift assay (B) detected by western blot (C), and efficiency quantification (D) (n=5). (E,F) Schematic, imaging (E) and quantification (F) of endothelial transwell migration when exposed to varying concentrations of Az-VEGF or non-Az-VEGF. (G) Quantification of endothelial metabolism when exposed to varying concentrations of Az-VEGF or non-Az-VEGF. For E-H, cells treated with complete EGM-2 served as positive controls (n=3) and cells treated with EBM-2 served as negative control. One-way ANOVA was used for statistical analysis and quantitative data were displayed as with mean <u>±</u> SEM, * *p*<0.05, ** *p*< 0.01, *** *p*<0.001.

Azido labeling of VEGF165 was achieved by leveraging the inherent cellular metabolic pathway for post-translational modification of newly synthesized proteins, particularly the sialic acid glycosylation [29, 30]. Mammalian Chinese hamster ovary (CHO) cells were used as the cell source, as they allow for recombinant proteins to be expressed in their glycosylated form [31, 32], and were transduced with a lentiviral vector expressing human VEGF165. For metabolic glycan labeling, VEGF-expressing CHO cells were cultured for 72 hours with 50 µM of the azide-functionalized Ac_4_ManNAz, an analog of the sialic acid biosynthetic precursor, N-acetylmannosamine (ManNAc). For the control culture to produce VEGF165 without azido labeling (non-Az-VEGF), cells were treated with 0.1% DMSO (vehicle) for the same duration. Following the labeling period, conditioned medium was collected from the CHO cell culture, and recombinant VEGF165 was affinity-purified through its C-terminal His tag.

To assess azido tag incorporation in the recombinant VEGF165, a click shift assay was performed involving the protein of interest conjugated with DBCO-PEG (30 kDa) and analyzed by Western blot. Only the azido-tagged protein will lead to successful DBCO-PEG conjugation and consequently a shift (increase) in molecular weight (Figure 1B). The click shift assay demonstrated over 50% azido labeling efficiency of VEGF165 from the Az-VEGF culture (Figure 1C lane 1), while no visible azido labeling was observed in the control non-Az-VEGF culture (Figure 1C lanes 2-4, Figure 1D, Figure S1). To assess and compare the bioactivity of Az-VEGF and non-Az-VEGF, we performed a VEGF-stimulated chemotactic endothelial transwell migration assay (Figure 1E,F) and an endothelial cell metabolic assay (Figure 1G), finding that both VEGF species, with and without azido labeling, induced comparable endothelial responses in a dose-dependent manner. Taken together, our data successfully demonstrated azido labeling of VEGF165 without affecting its bioactivity.

### 3.2 Metabolic glycan engineering of eGFP that is natively non-glycosylated

Despite the popularity of glycosylated GFs, there are other GFs and proteins of interest for tissue engineering and regeneration that are not natively glycosylated in their mature forms [33-35]. With this in mind, here we intend to extend the utility of our chemoselective, site-specific azido tagging platform to these natively non-glycosylated proteins, using eGFP, as a model protein for demonstration. For glycosylation to take place, the newly synthesized protein requires two critical components: a signal peptide (SP) that directs new protein synthesis to the endoplasmic reticulum (ER) and Golgi apparatus; and a glycosylation sequon that serves as the attachment site for oligosaccharide, frequently an N-linked glycan. For VEGF165, both the signal peptide (MNFLLSWVHWSLALLLYLHHAKWSQA) and sequon (SNITM) have been identified [36]. Here, we intend to convert the natively non-glycosylated eGFP into an N-linked glycoprotein for site-specific azido incorporation by engineering its amino acid sequence to include the VEGF165-derived signal peptide (SP_VEGF_) at its N-terminus and three tandem repeats of the VEGF165-derived sequon (Sequon_VEGF_) at its C-terminus, resulting in SP_VEGF_-eGFP-Sequon_VEGF_ (Figure 2A, Table S1). For comparison, we also created expression vectors for unmodified eGFP and eGFP with only the sequon but not the signal peptide (referred to as eGFP-Sequon_VEGF_).

**Figure 2.**
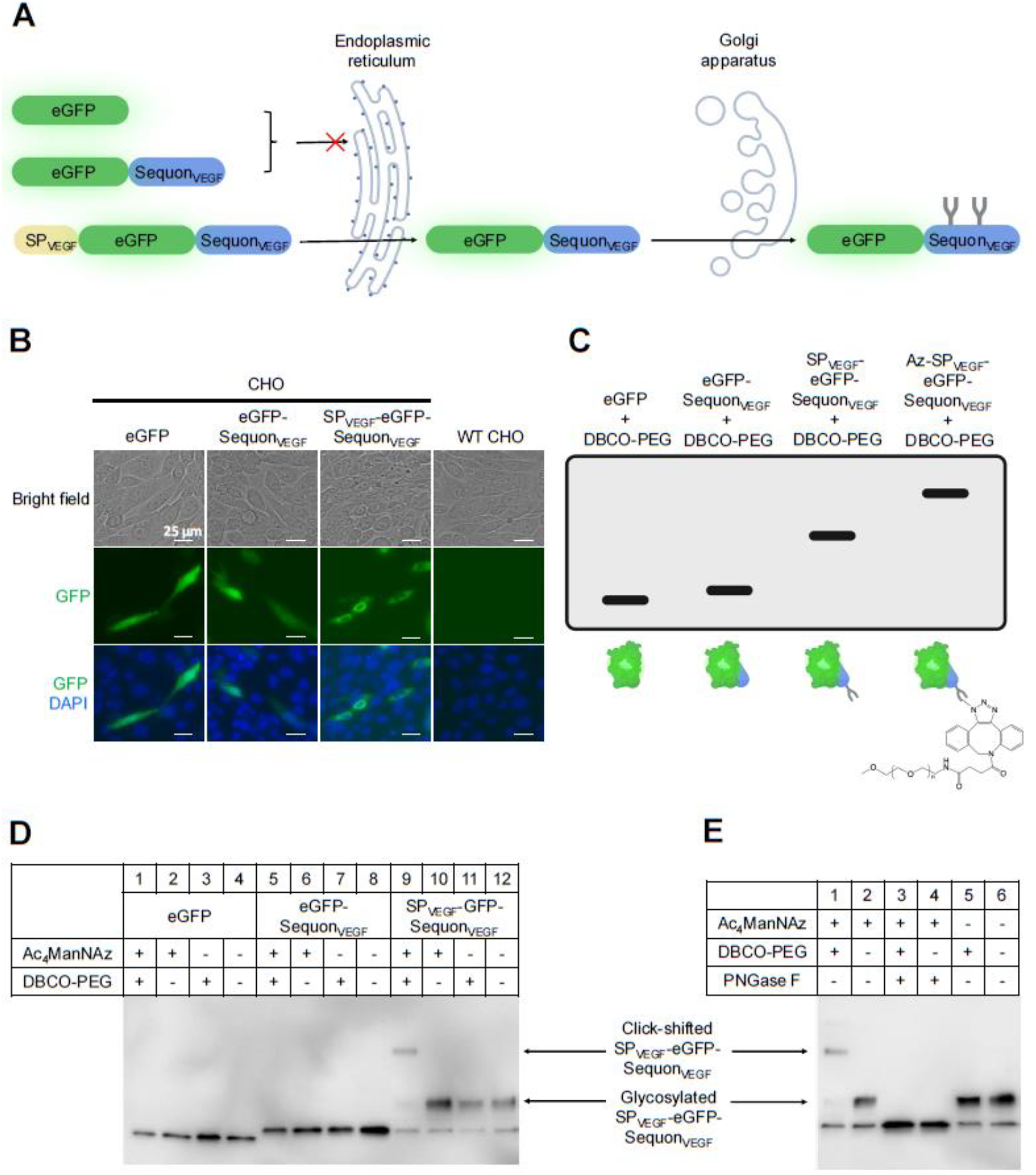
Metabolic glycan engineering of eGFP. (A) Schematic showing the overall strategy for glycan engineering of eGFP. Created with BioRender.com. (B) Sub-cellular localization of CHO cells expressing native and engineered eGFP. (C) Schematic showing click-shift assay of native and engineered eGFP through conjugation. (D) Western blot analysis of click-shift assay for native and engineered eGFP with and without metabolic glycan labeling. (E) Western blot analysis of click-shift assay following deglycosylation with PNGase F for SP_VEGF_-eGFP-Sequon_VEGF_.

After transfection with all eGFP vectors (both native and engineered) in CHO cells, we initially investigated the subcellular localization of eGFP by utilizing its inherent fluorescence. While the unmodified eGFP and eGFP-Sequon_VEGF_ exhibited widespread fluorescence throughout the cytoplasm, the fluorescence signal in cells transfected with SP_VEGF_-eGFP-Sequon_VEGF_ was confined to the perinuclear region. This suggests that the inclusion of the VEGF-derived signal peptide successfully redirected SP_VEGF_-eGFP-Sequon_VEGF_ into the protein secretory pathway essential for glycosylation to occur (Figure 2B). In agreement with this, in gel electrophoresis, only SP_VEGF_-eGFP-Sequon_VEGF_ displayed a higher molecular-weight band (Figure 2C, 2D lanes 9-12), corresponding to glycosylated eGFP, which was absent in WT-eGFP or eGFP-Sequon_VEGF_ (Figure 2C, 2D lanes 1-8). Moreover, a click shift assay using DBCO-PEG conjugation resulted only in an upward molecular weight shift of SP_VEGF_-eGFP-Sequon_VEGF_ produced by cells pre-treated with Ac_4_ManNAz (Figure 2C, 2D lane 9), further supporting the successful glycosylation and azido incorporation of SP_VEGF_-eGFP-Sequon_VEGF_. To further validate this and specifically examine the site specificity of azido incorporation during glycosylation, the Ac_4_ManNAz-labeled SP_VEGF_-eGFP-Sequon_VEGF_ was digested with PNGase F, which cleaves between the innermost GlcNAc and asparagine residue in N-linked oligosaccharides, and then subjected to the click shift assay. PNGase F-mediated deglycosylation completely eliminated the bands corresponding to glycosylated SP_VEGF_-eGFP-Sequon_VEGF_ and its PEG conjugate (Figure 2E lanes 3-4), consistent with the complete removal of N-linked oligosaccharides from glycoproteins by PNGase F. This confirms that Ac_4_ManNAz-mediated azido incorporation specifically targets the glycan modification of the engineered SP_VEGF_-eGFP-Sequon_VEGF_ glycoprotein.

We observed similar subcellular localization and glycosylation-dependent azido incorporation when we switched to an alternative signal peptide from the rat Follicle-Stimulating Hormone β-subunit [22] (Figure S2A,B). Taken together, our data demonstrate that non-glycosylated eGFP can be engineered to undergo glycosylation through the incorporation of a signal peptide and sequon, further enabling site-specific azido incorporation on the glycan chain of the newly synthesized SP-eGFP-Sequon glycoprotein. This illustrates the generalizability of our metabolic glycan engineering strategy to be applied to both natively glycosylated and non-glycosylated proteins.

### 3.3 Engineering and characterization of azide-reactive ECM hydrogel

Fibrin hydrogel is a widely used ECM biomaterial for tissue engineering and regenerative medicine applications [7, 37]. Using amine-reactive carbodiimide chemistry [38], we engineered DBCO-conjugated, azide-reactive fibrinogen (DBCO-fibrinogen) for fibrin hydrogel formation and chemoselective immobilization of Az-VEGF. To enable effective DBCO-NHS reaction with primary amines on fibrinogen while maintaining ECM solubility (preventing polymerization), the reaction was performed with diluted fibrinogen (10 mg/mL) in PBS (Figure 3A). To assess DBCO modification, the resulting DBCO- and unmodified control fibrinogen was incubated with Az-Cy5, and subsequent dot blot analysis showed robust and specific Cy5 conjugation to DBCO-fibrinogen (Figure 3B), demonstrating effective DBCO modification of fibrinogen.

**Figure 3.**
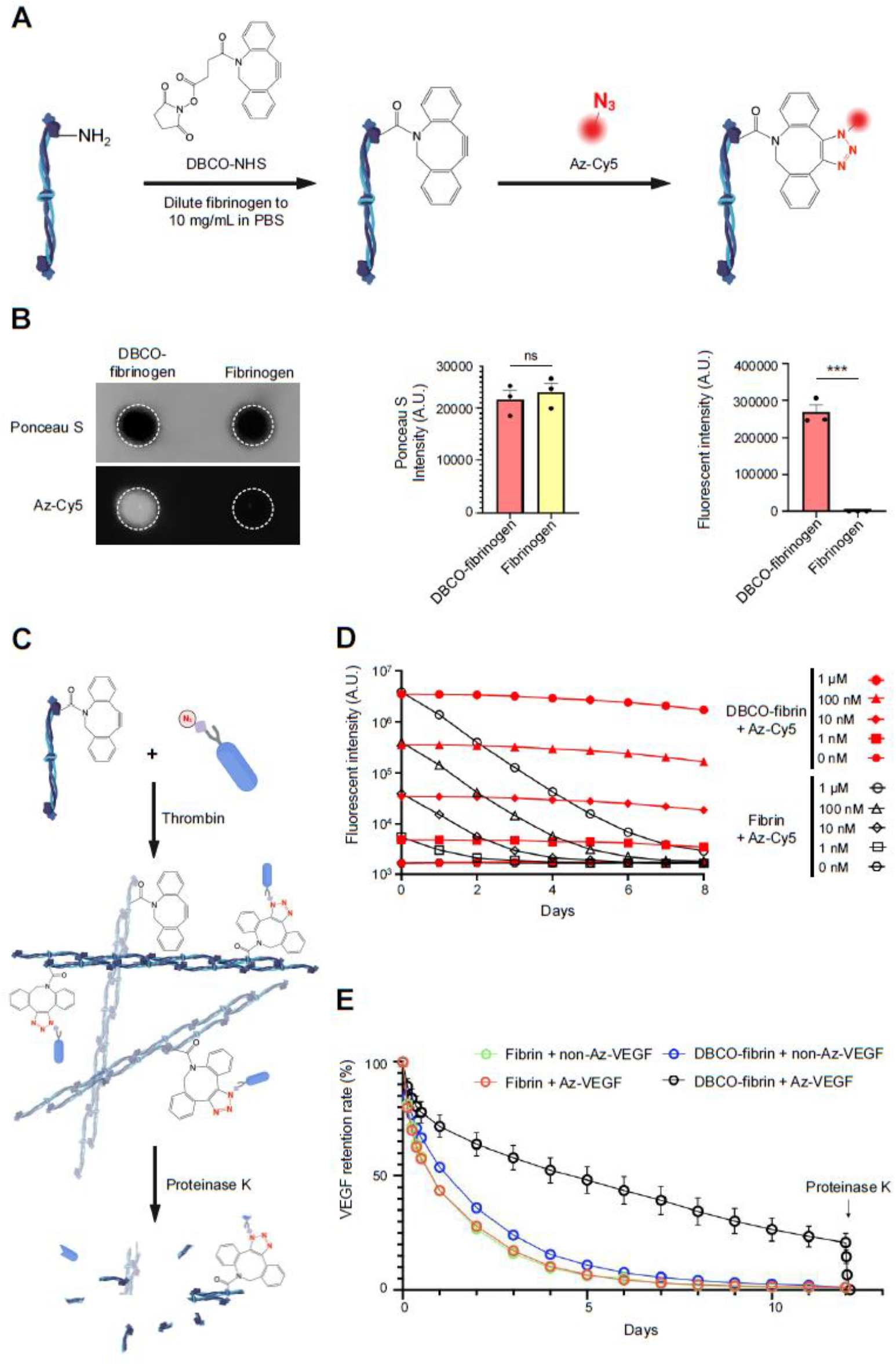
Engineering and characterization of azide-reactive DBCO-fibrin hydrogel. (A) Schematic showing production of DBCO-fibrinogen via the conjugation between primary amines (−NH_2_) on fibrinogen and DBCO-NHS, and subsequent click-conjugation between DBCO-fibrinogen and Az-Cy5. Created partly with BioRender.com. (B) Dot blot analysis and quantification of DBCO modification of fibrinogen following click-conjugation with Az-Cy5, equal protein loading was confirmed by Ponceau S staining (n=3). (C) Schematic showing DBCO-fibrinogen click-conjugation with Az-labeled molecules, followed by fibrin hydrogel formation and proteinase K digestion. (D) Quantification of fluorescence retention of Az-Cy5 in DBCO-fibrin and unmodified fibrin hydrogels (n=3). (E) Quantification of the retention of radiolabeled Az-VEGF and non-Az-VEGF in DBCO-fibrin and unmodified fibrin hydrogels (n=3). Student’s t-test was used for panel B while one-way ANOVA was used for panel D and E, and quantitative data were displayed as with mean <u>±</u> SEM, * *p*<0.05, ** *p*< 0.01,*** *p*<0.001.

To evaluate the ability of DBCO-modified fibrin (DBCO-fibrin) gel to retain azido molecules long-term, fibrin hydrogels were created from either DBCO- or control fibrinogen (15 mg/mL) and assessed for the retention of various concentrations of Az-Cy5. While the fluorescence of Az-Cy5 decreased rapidly when incorporated into unmodified fibrin hydrogel, it remained nearly constant in DBCO-fibrin for over 8 days (Figure 3C,D, Table S2). Meanwhile, a clear dose-dependence in Az-Cy5 fluorescence retention can be observed, and click-mediated enhancement in Az-Cy5 retention within DBCO-fibrin can be detected with input concentrations as low as 1 nM of Az-Cy5, demonstrating the high sensitivity of DBCO-fibrin for the immobilization of azido molecules.

We examined whether similar specificity within DBCO-fibrin hydrogel can be achieved for the click-immobilization of VEGF165 with site-specific azido incorporation. ^125^I-radiolabeled Az-VEGF and non-Az-VEGF were introduced to DBCO-fibrin and control hydrogels, and their retention was monitored. The combination of ^125^I-Az-VEGF and DBCO-fibrin resulted in superior long-term retention of VEGF radioactivity, over 17 times higher than all other ^125^I-VEGF-fibrin combinations (Figure 3E, Table S3). To assess the biodegradability of the click-immobilized VEGF within the fibrin hydrogel, we introduced proteinase K on Day 12 and observed the immediate release of over 99.8% of the VEGF retained in DBCO-fibrin (Figure 3E). Overall, these results demonstrate that glycan-engineered VEGF with site-specific azido modification can be effectively immobilized within DBCO-modified ECM hydrogels with robust long-term retention.

### 3.4 Evaluation of angiogenic functions of VEGF-immobilized hydrogel

To verify the functionality of the click-immobilized VEGF within DBCO-fibrin hydrogel, we conducted a series of in vitro endothelial assays using HUVECs as the model. In these assays, we prepared various hydrogel conditions with combinations of DBCO- (or control unmodified) fibrin hydrogel and Az-VEGF (or control non-Az-VEGF) at different doses.

First, we conducted an endothelial metabolic assay using a double-layer hydrogel system, where the pre-washed bottom-layer hydrogel contained VEGF-immobilized fibrin as the source, and the top-layer hydrogel embedded with HUVECs to detect and respond to the released VEGF (Figure 4A). The metabolic activity of the embedded HUVECs was measured using the resazurin metabolism assay, which demonstrated a significant upregulation of top-layer endothelial metabolic activity when cells were exposed to the bottom-layer hydrogel of DBCO-fibrin with 200 ng/mL Az-VEGF compared to all other conditions that failed to achieve click-immobilization of the introduced VEGF (48% higher than DBCO-fibrin with 200 ng/mL non-Az-VEGF) (Figure 4B). Additionally, the same hydrogel combinations were used for the endothelial scratch recovery assay and the transwell endothelial chemotaxis assay, where the hydrogel acted as the source of sustained VEGF release to promote endothelial migration. In the scratch recovery assay, the hydrogel was placed in the upper chamber of the transwell with HUVECs in the bottom chamber, while the arrangement was reversed for the transwell chemotaxis assay (Figures 4A). The scratch recovery in the HUVEC monolayer was expressed as the percentage of scratch closure over 24 h post-scratch induction, and the coverage of migrating HUVECs was quantified after 24 h of culture. The DBCO-fibrin hydrogel with 200 ng/mL Az-VEGF immobilization showed 50% higher scratch closure in the scratch recovery assay and induced two-fold higher HUVEC migration in the transwell chemotaxis assay compared to non-clicked DBCO-fibrin with 200 ng/mL non-Az-VEGF (Figure 4C-F). When compared with the groups of 50 ng/mL Az-VEGF-immobilized DBCO-fibrin, the dose-dependent angiogenic responses of HUVECs were observed in all three assays. Together, as a source for sustained VEGF release, VEGF-click-immobilized fibrin hydrogels demonstrated enhanced pro-angiogenic properties.

**Figure 4.**
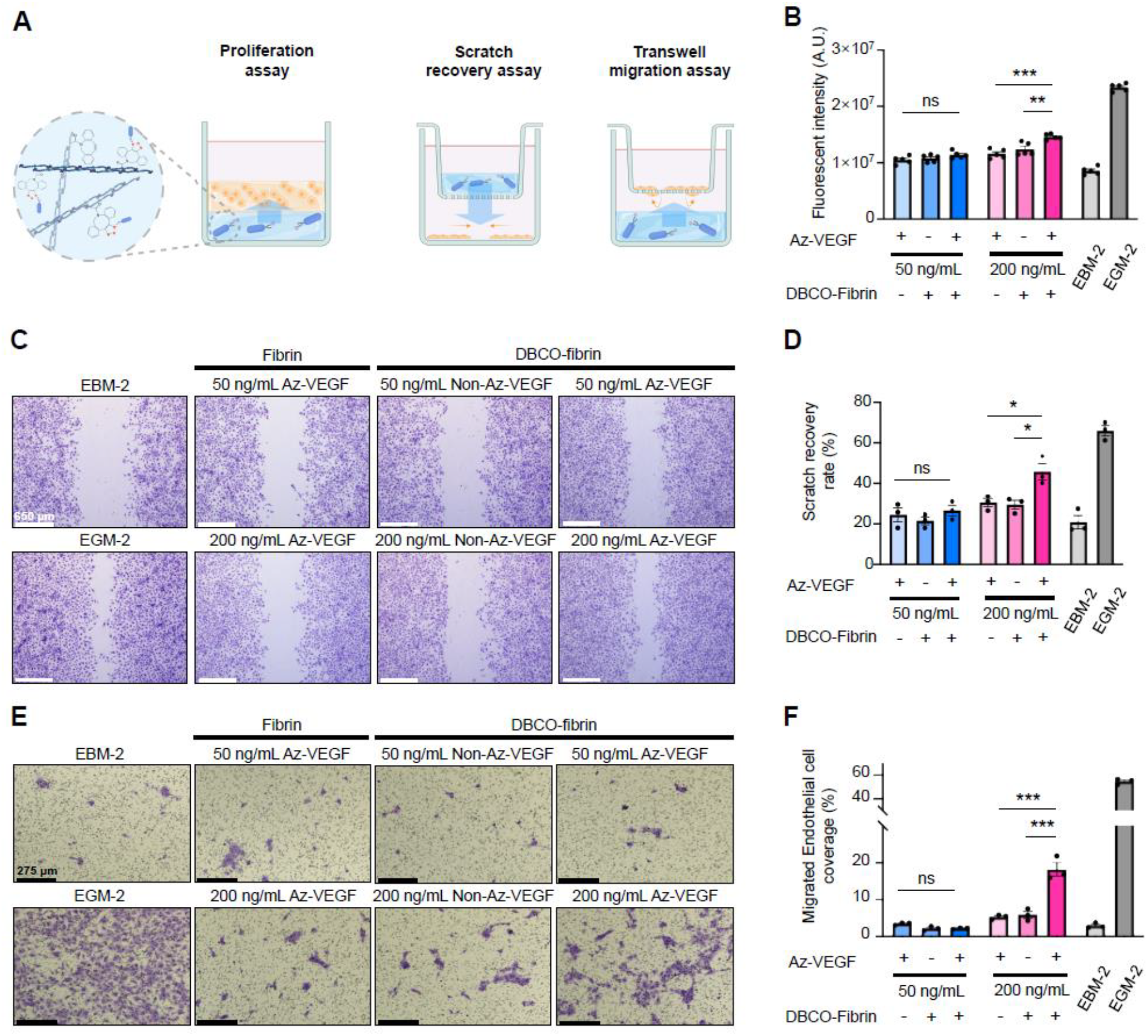
Evaluation of angiogenic functions of VEGF-immobilized hydrogel. (A) Schematic showing double-layer fibrin gel for metabolic assay with HUVECs 3D culture (left) and endothelial transwell migration (right) when exposed to fibrin gels with varying concentrations of Az-VEGF or non-Az-VEGF. (B) Quantification of endothelial metabolism in double-layer fibrin gel with varying concentrations of Az-VEGF or non-Az-VEGF. (C,D) Imaging (C) and quantification (D) of endothelial scratch recovery assay when exposed to fibrin gels with varying concentrations of Az-VEGF or non-Az-VEGF. (E,F) Quantification (E) and imaging (F) of endothelial transwell migration assay when exposed to fibrin gels with varying concentrations of Az-VEGF or non-Az-VEGF. Cells treated with complete EGM-2 with unmodified fibrin gel served as positive controls (n=3) and cells treated with EBM-2 with unmodified fibrin gel served as negative control. One-way ANOVA was used for statistical analysis and quantitative data were displayed as with mean ± SEM, * *p*<0.05, ** *p*< 0.01, *** *p*<0.001.

## 4. Discussion

In this study, we present a strategy for site-specific glycan engineering of GFs to incorporate click chemistry-reactive azido tags. This approach is applicable to the recombinant expression of glycoproteins (such as VEGF165) and can be extended through engineered glycosylation to modify proteins that are non-glycosylated in their natural forms (such as eGFP). Additionally, we demonstrated that VEGF165 with site-specific azido tagging can be effectively immobilized within ECM hydrogel containing the complementary DBCO modification through copper-free click conjugation, leading to sustained release and enhanced pro-angiogenic functions.

In native tissue environments, GFs often physically associate with the ECM framework by binding to fibrous proteins (such as collagens) or glycosaminoglycans (like heparan sulfate), which regulates the bioavailability, stability, and signaling dynamics of GFs[7, 39]. This observation has inspired extensive efforts in biomaterial engineering aimed at enhancing GF functionalities by recapitulating or even augmenting their association with ECM materials, with the goal of preventing rapid diffusion and degradation of GFs to ensure localized and sustained biological functions. While numerous strategies have been developed to promote GF-ECM binding, most of these approaches require individual optimizations due to the distinct biochemical natures of the GFs and ECM materials being investigated. The approach described here overcomes the limitations of conventional protein conjugation by providing a platform technology applicable to nearly all recombinant proteins, through site-specific incorporation of azido tags on native or engineered glycosylation. This enables subsequent biomaterial conjugation to be conducted via click chemistry with high specificity and efficiency.

While glycosylated GFs have been well-documented for their critical contributions to tissue engineering, there are also non-glycosylated GFs, such as fibroblast growth factor-2, epidermal growth factor, and insulin-like growth factor-1, that are extensively used in stem cell and tissue engineering [40-44]. To broaden the applicability of our platform to non-glycosylated proteins, using eGFP as an example, we engineered its sequence to include a signal peptide that directs new protein synthesis to the endoplasmic reticulum-Golgi secretory pathway, where glycosylation occurs, along with glycosylation sequons that serve as attachment sites for polysaccharides [22, 45]. The inclusion of these two elements effectively altered the subcellular localization of the engineered eGFP and enabled its subsequent azido tagging, which was sensitive to PNGase digestion, suggesting glycan-specific azido incorporation. This demonstrates that our GF glycan engineering can be adapted to a broader range of proteins, including those that are not natively glycosylated. With the bioorthogonally incorporated click moieties, the glycan-engineered proteins can be conveniently conjugated with other biomolecules bearing complementary alkyne or DBCO groups. Beyond ECM hydrogel functionalization demonstrated here, the utility of the presented technology can be extended to the engineering of drug-conjugates of proteins, such as antibodies, that are commonly glycosylated. In this study, we showed that the engineered eGFP exhibited even higher azido labeling efficiency than the native VEGF165, which can be explained by the increase in the number of sequons from one in VEGF165 to three in the engineered eGFP. This also provides a potential approach to enhance metabolic glycan engineering efficiency by incorporating tandem repeats of sequons.

Our strategy leverages the intrinsic ability of mammalian cells to perform post-translational glycosylation, a natural biosynthetic process that ensures recombinant proteins like VEGF165 are naturally modified with glycan chains during production [46-48]. Although protein glycosylation has been demonstrated to protect proteins from proteolytic degradation and rapid clearance [49], it is also generally considered dispensable for the fundamental biological functions of these proteins [50, 51]. For example, while bacterial cells do not carry out post-translational glycosylation as eukaryotic cells do [52], bacterial expression systems have been widely utilized for producing functional recombinant proteins, many of which are GFs [53-56]. Here, we build on these concepts regarding protein glycosylation and further enhance glycosylation-based modulation of protein stability by introducing chemoselective ligands on the recombinant GFs for biomaterial conjugation, thereby boosting GF long-term retention and function.

Beyond Ac_4_ManNAz, which is utilized here for targeting protein sialylation, N-azidoacetylgalactosamine (GalNAz) and N-azidoacetylglucosamine (GlcNAz) are also widely employed in metabolic glycan engineering within living systems [57], offering the potential for incorporating azido-labeling into the recombinant expression of proteins bearing other types of glycans, such as mucin-type O-linked glycans [58-60]. In addition to mammalian expression systems, bacteria are also popular host cells for recombinant protein expression. While native bacteria generally cannot produce glycoproteins with complex, eukaryotic-like glycosylation patterns, efforts are underway to introduce foreign glycotransferases to enable mammalian-type glycosylation in engineered bacteria such as E. coli [61, 62]. This expands the utility of the metabolic glycan engineering strategy discussed here to include recombinant glycoprotein expression in engineered bacterial strains capable of performing mammalian-like glycosylation.

In summary, we have developed a versatile platform for the metabolic glycan engineering of GFs and the chemoselective immobilization of engineered GFs onto ECM-derived hydrogels. This platform allows GF engineering without relying on individually designed approaches. The ECM functionalization strategy established in our studies is highly amenable to modern 3D biofabrication techniques, such as bioprinting, enabling the creation of biomaterials and tissue constructs with spatially confined presentation of GFs and other morphogenic proteins to enhance the precision of tissue pattern registration [63, 64].

## 5. Conclusion

In this study, we developed a novel GF engineering platform that metabolically alters the glycans of GFs using azido tags and immobilizes them within a fibrin hydrogel through chemoselective click chemistry. This approach enhances GF retention and promotes in vitro angiogenic responses in endothelial cells.

## Author Contributions

Y.X., Q.L., P.G.C. and X.R. contributed to the project design, data interpretation and manuscript preparation.

Y.X. and Q.L. performed the experiments and analyzed data.

## Acknowledgement

This work was supported by Shenzhen Excellent Technology Co., Ltd, and the Department of Biomedical Engineering at Carnegie Mellon University. Y.X. was partly supported by scholarships from the China Scholarship Council. We are grateful to Misti West and Garrett Struble for laboratory management.

## Data Availability

The authors declare that all data supporting the findings of this study are available within the article and its supplementary material files, or from the corresponding author on reasonable request.

