## Supplemental materials for "Glycosylation-enabled Site-specific Growth Factor Engineering for Biomaterial Functionalization"

### Supplementary Figures

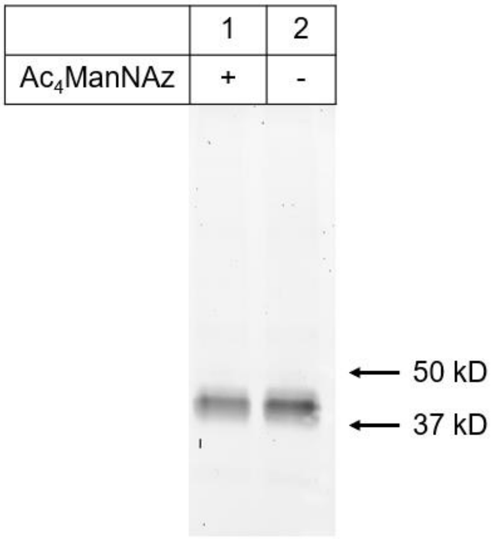


**Fig S1. Sypro Ruby protein gel staining of purified Az-VEGF and non-Az-VEGF.**

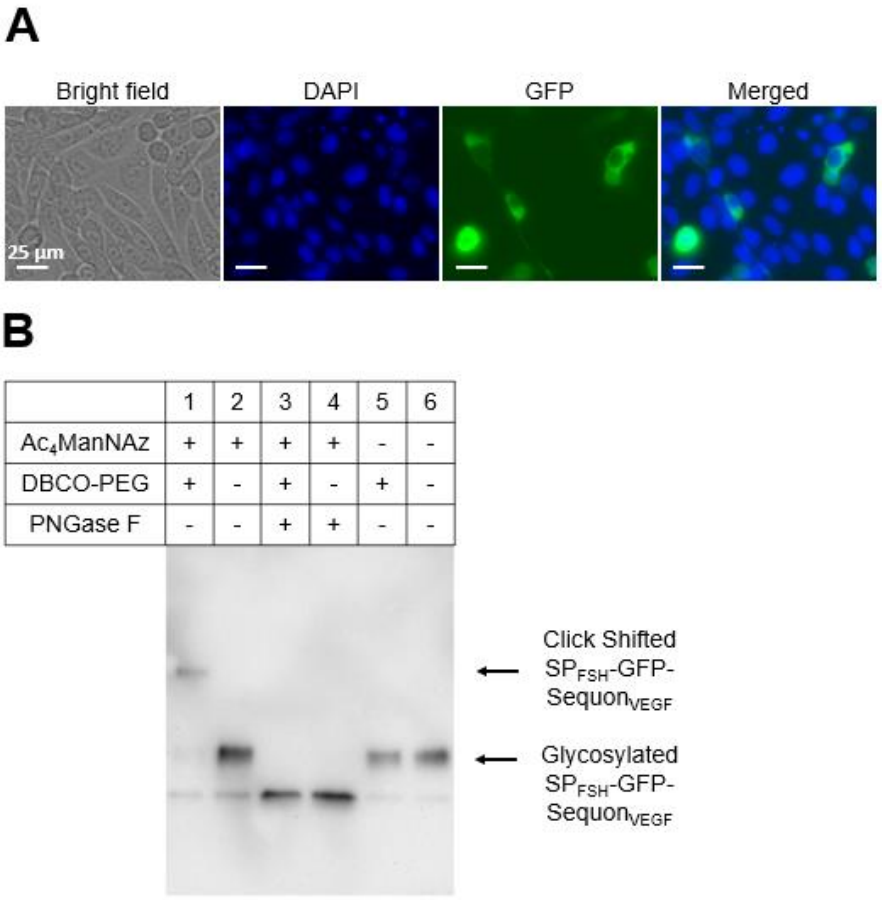


**Fig S2. Metabolic glycan engineering of eGFP with $SP_{FSH}$.** (A) Subcellular localization of $SP_{FSH}$-eGFP in CHO cells. (B) Western blot analysis of click-shift assay of $SP_{FSH}$-eGFP-$Sequon_{VEGF}$ with and without metabolic glycan labeling, and with and without deglycosylation with PNGase F.

### Supplementary Tables

| Vector | AA sequence |
|---|---|
| VEGF165 | MNFLLSWVHWSLALLLYLHHAKWSQAAPMAEGGGQNHHEVVKFMDVYQRSYCHP<br>IETLVDIFQEYPDEIEYIFKPSCVPLMRCGGCCNDEGLECVPTEESNITMQIMRIKPH<br>QGQHIGEMSFLQHNKCECRPKKDRARQENPCGPCSERRKHLFVQDPQTCKCSCK<br>NTDSRCKARQLELNERTCRCDKPRRGGGGSENLYFQGHHHHHH* |
| eGFP | MVSKGEELFTGVVPILVELDGDVNGHKFSVSGEGEGDATYGKLTLKFICTTGKLPVP<br>WPTLVTTLTYGVQCFSRYPDHMKQHDFFKSAMPEGYVQERTIFFKDDGNYKTRAE<br>VKFEGDTLVNRIELKGIDFKEDGNILGHKLEYNYNSHNVYIMADKQKNGIKVNFKIRH<br>NIEDGSVQLADHYQQNTPIGDGPVLLPDNHYLSTQSALSK DPNEKRDHMVLLEFVT<br>AAGITLGMDELYK* |
| eGFP-Sequon$_{VEGF}$ | MVSKGEELFTGVVPILVELDGDVNGHKFSVSGEGEGDATYGKLTLKFICTTGKLPVP<br>WPTLVTTLTYGVQCFSRYPDHMKQHDFFKSAMPEGYVQERTIFFKDDGNYKTRAE<br>VKFEGDTLVNRIELKGIDFKEDGNILGHKLEYNYNSHNVYIMADKQKNGIKVNFKIRH<br>NIEDGSVQLADHYQQNTPIGDGPVLLPDNHYLSTQSALSK DPNEKRDHMVLLEFVT<br>AAGITLGMDELYKGGGGSSNITMSNITMSNITMENLYFQGHHHHHH* |
| SP$_{VEGF}$-eGFP-Sequon$_{VEGF}$ | MNFLLSWVHWSLALLLYLHHAKWSQAVSKGEELFTGVVPILVELDGDVNGHKFSVS<br>GEGEGDATYGKLTLKFICTTGKLPVPWPTLVTTLTYGVQCFSRYPDHMKQHDFFKS<br>AMPEGYVQERTIFFKDDGNYKTRAEVKFEGDTLVNRIELKGIDFKEDGNILGHKLEY<br>NYNSHNVYIMADKQKNGIKVNFKIRHNIEDGSVQLADHYQQNTPIGDGPVLLPDNHY<br>LSTQSALSK DPNEKRDHMVLLEFVTAAGITLGMDELYKGGGGSSNITMSNITMSNIT<br>MENLYFQGHHHHHH* |
| SP$_{FSH}$-eGFP-Sequon$_{VEGF}$ | MMKSIQLCILWCLRAVCCHVSKGEELFTGVVPILVELDGDVNGHKFSVSGEGEGDA<br>TYGKLTLKFICTTGKLPVPWPTLVTTLTYGVQCFSRYPDHMKQHDFFKSAMPEGYV<br>QERTIFFKDDGNYKTRAEVKFEGDTLVNRIELKGIDFKEDGNILGHKLEYNYNSHNV<br>YIMADKQKNGIKVNFKIRHNIEDGSVQLADHYQQNTPIGDGPVLLPDNHYLSTQSAL<br>SK DPNEKRDHMVLLEFVTAAGITLGMDELYKGGGGSSNITMSNITMSNITMENLYF<br>QGHHHHHH* |

**Table S1. Amino acid sequences of VEGF165, eGFP, eGFP-Sequon$_{VEGF}$, SP$_{VEGF}$-eGFP-Sequon$_{VEGF}$ and SP$_{FSH}$-eGFP-Sequon$_{VEGF}$.**

| Fibrin | Cy5 | 1 μM | 100 nM | 10 nM | 1nM | 0nM |
|---|---|---|---|---|---|---|
| Days | 0 | *** | * | *** | ** | ns |
| | 1 | *** | *** | *** | *** | ns |
| | 2 | *** | *** | *** | *** | ns |
| | 3 | *** | *** | *** | *** | ns |
| | 4 | *** | *** | *** | *** | ns |
| | 5 | *** | *** | *** | *** | ns |
| | 6 | *** | *** | *** | *** | ns |
| | 7 | *** | *** | *** | *** | ns |
| | 8 | *** | *** | *** | *** | ns |
| | 9 | *** | *** | *** | *** | ns |
| | 10 | *** | *** | *** | *** | ns |

**Table S2. Summary of *p* values from comparisons of Cy5 fluorescence retention kinetics of DBCO-fibrin vs. unmodified fibrin hydrogels at varying initial Az-Cy5 concentrations. * $p<0.05$, ** $p< 0.01$, *** $p<0.001$.**

| | | Fib | Fib | DBCO-Fib |
|---|---|---|---|---|
| | VEGF | Az | NA | NA |
| Times | 0h | ns | ns | ns |
| | 3h | * | ns | ns |
| | 6h | ** | ** | ns |
| | 9h | ** | ** | ns |
| | 12 h | ** | ** | * |
| | 24 h | *** | *** | ** |
| | Day 2 | *** | *** | *** |
| | Day 3 | *** | *** | *** |
| | Day 4 | *** | *** | *** |
| | Day 5 | *** | *** | *** |
| | Day 6 | *** | *** | *** |
| | Day 7 | *** | *** | *** |
| | Day 8 | *** | *** | *** |
| | Day 9 | *** | *** | *** |
| | Day 10 | *** | *** | *** |
| | Day 12 | *** | *** | *** |

**Table S3. Summary of *p* values from the comparisons of Az-VEGF and non-Az-VEGF retention kinetics in DBCO-fibrin or unmodified fibrin hydrogels. * $p<0.05$, ** $p< 0.01$, *** $p<0.001$.**
